# CaDrA: A computational framework for performing candidate driver analyses using binary genomic features

**DOI:** 10.1101/221846

**Authors:** Vinay K. Kartha, Joseph G. Kern, Paola Sebastiani, Liye Zhang, Xaralabos Varelas, Stefano Monti

## Abstract

Identifying complementary genetic drivers of a given phenotypic outcome is a challenging task that is important to gaining new biological insight and discovering targets for disease therapy. Existing methods aimed at achieving this task lack analytical flexibility. We developed Candidate Driver Analysis or CaDrA, a framework to identify functionally-relevant subsets of binary genomic features that, together, are associated with a specific outcome of interest. We evaluate CaDrA’s sensitivity and specificity for typically-sized multi-omic datasets, and demonstrate CaDrA’s ability to identify both known and novel drivers of oncogenic activity in cancer cell lines and primary tumors.

## Background

Advances in high-throughput sequencing technology has led to a rapid rise in the availability of large multi-omic datasets through compendia such as the Cancer Cell Line Encyclopedia (CCLE), The Cancer Genome Atlas (TCGA), the Genotype-Tissue Expression (GTEx), and others [1–3]. These data include genetic alterations, comprising somatic copy number alterations (SCNAs) and somatic mutations, epigenetic information, such as microRNA expression and DNA methylation, as well as gene expression profiling through microarray or RNA-sequencing (RNASeq) technology, across tens of thousands of samples representing varying biological contexts. Concomitantly, several computational methods have been developed and applied to effectively query and integrate different types of genome-wide datasets in order to make meaningful predictions about the biological processes driving the phenotypes of interest [4,5]. An important application of such methods is the identification of recurrent genomic alterations, and their potential effects on downstream pathway activity or phenotypes. For example, in many cancers, samples exhibiting elevated activity of a given oncogenic signature may be enriched for, or driven by functionally-relevant somatic mutations or SCNAs. Identifying such associations may help unravel underlying mechanisms contributing to abnormal pathway activity, further enabling disease subtyping and sample classification [6–8]. Alternatively, linking these genomic features with phenotypic readouts such as drug sensitivity may support the discovery of novel druggable targets and further guide precision medicine regimens [9–11].

Recently, computational methods have been developed to identify subsets or combinations of genomic features that are collectively associated with a given phenotypic response [12–15]. These methods, while having the capacity to find alternative drivers contributing to the same downstream effect, tend to rely on restrictive properties such as mutual exclusivity of features, or feature recurrence in a subset of samples independent of the target phenotypic profile under consideration. More importantly, not all methods support the joint analyses of features including SCNAs and somatic mutations, with possible extension to other genomic data, while also allowing for multiple statistical feature scoring functions, and rigorous assessment of the statistical significance of the discovered associations. Finally, a user-friendly and flexible programming package supporting the rapid screening for candidate drivers given a set of ranked genomic features is currently lacking, and would prove extremely useful for incorporation in analytical pipeline frameworks aimed at the generation of novel biological hypotheses.

Here, we present Candidate Driver Analysis (CaDrA), a methodology that searches for the set of genomic alterations, here denoted as *features* (mutations, somatic copy number alterations, translocations, etc.) associated with a user-provided ranking of samples within a dataset. Our method specifically employs a stepwise heuristic search to identify a subset of features whose union is maximally-associated with the observed sample ranking, and carries out rigorous statistical significance testing based on sample permutation, thereby allowing for the identification of candidate genetic drivers associated with aberrant pathway activity or drug sensitivity, while still exploiting aspects of feature complementarity and sample heterogeneity. To highlight the method’s overall performance, along with its relevance and ability to select sets of genomic features that indeed drive certain oncogenic phenotypes in cancer, we apply CaDrA to simulated data, as well as to real genomic data corresponding to cancer cell lines and primary human tumors. The results from simulations suggest CaDrA displays high sensitivity for mid- to large-sized datasets, and high specificity for all sample sizes considered. Using genomic data drawn from CCLE and TCGA, we demonstrate CaDrA’s capacity to correctly identify well-characterized driver mutations in cancer cell lines, along with its ability to discover less-known features associated with invasive phenotypes in human breast cancer samples, which we functionally validate *in vitro*. Our package, which is publicly-available, will allow for rapidly mining numerous multi-omics datasets for candidate drivers of user-specified molecular readouts, such as pathway activity, drug sensitivity, protein expression, or other quantitative measurements of interest, further enabling targeted queries and novel hypothesis generation.

## Results

### CaDrA overview

An overview of CaDrA’s workflow is summarized in Figure 1. CaDrA implements a step-wise heuristic approach that searches through a set of binary features (each represented as a 1/0-valued vector, indicating the presence/absence of a SCNA, somatic mutation, or other (epi)genetic alterations across samples, respectively), and returns a final subset of features whose union (logical OR) defines an alteration ‘metafeature’ that is maximally associated with the defined sample ranking provided as input (see Methods). The strength of the association of a meta-feature with a sample ranking is a function of the agreement between the skewness of the alterations’ occurrences and the sample ranking. The input sample ranking is usually a function of a sample-specific measurement, e.g., the activity level of a pathway, the response to a targeted treatment, the expression level of a given transcript or protein, etc. Therefore, the metafeature returned by the search is the set of features maximally predictive of that same sample-specific measurement variable. CaDrA allows for multiple modes to query ranked binary datasets with user-specified parameters defining search criteria, enables rigorous permutation-based significance testing of results, and reduced computation time by exploiting pre-computed score distributions and parallel computing, when available (see Methods).

**Figure 1.**
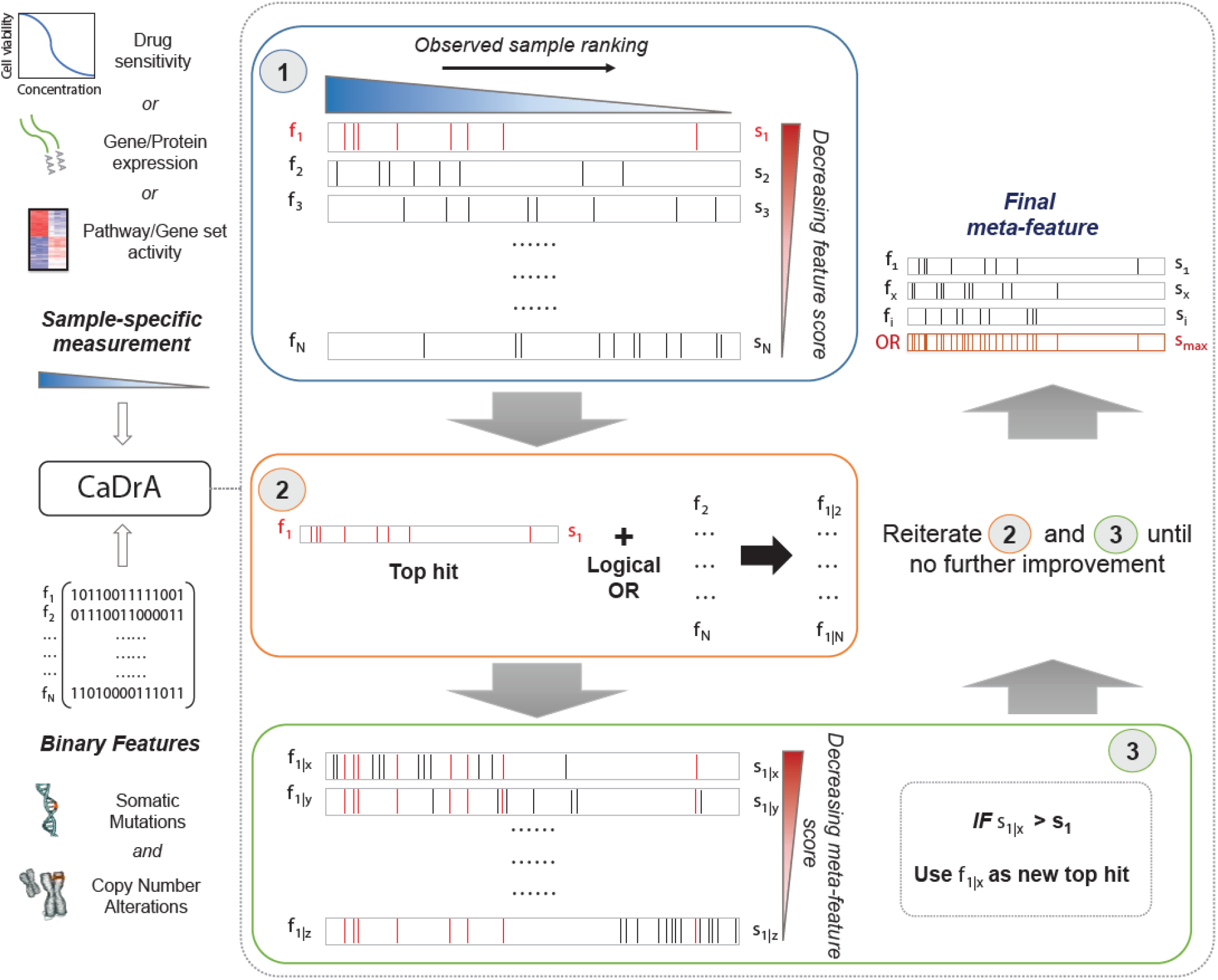
Overview of CaDrA workflow and implementation. CaDrA takes as input a sample-specific measurement to rank samples with, and a matrix of binary features of the same sample units. In Step 1 (blue box), CaDrA begins by choosing a starting feature, which is either the single feature having the best score based on its left-skewness, or a user-specified start feature. In the next step (Step 2; orange box), the union (logical OR) of this feature is taken with each of the remaining features in the dataset, yielding ‘meta-features’ with their corresponding scores. If any meta-feature has a better score than the hit from the previous step (Step 3; green box), CaDrA uses this new meta-feature as a reference for the next iteration, reiterating over Steps 2 and 3 until no further improvement in scores can be obtained. The final output is a set of features (meta-feature) whose union has the (local) maximal score.

### Analysis of simulated data to evaluate CaDrA performance

To assess the overall performance of CaDrA to recover (statistically) significantly associated meta-features, we simulated two types of datasets for a range of sample sizes: i) the *true-positive datasets* consist of both left-skewed (i.e. true positive with skewness concordant with sample ranking) as well as uniformly distributed (i.e. null) features; and ii) the *null datasets* consist of null features only (see Methods). This enabled us to estimate the overall sensitivity and specificity of CaDrA using the true positive and null datasets, respectively. By running CaDrA on multiple simulated datasets (*n*=500 true positive and null datasets, each), we first assessed the resulting meta-features based on the number of true positive features and the total number of features contained within each returned meta-feature (i.e., the meta-feature size; Fig. 2a and 2c). The true positive datasets had a maximum of 5 positive features to be detected, while the maximum number of features CaDrA was allowed to add was set to 7, to evaluate the ability of the search to recover all but no more than the positive features. The true positive rate (TPR) and false positive rate (FPR) of CaDrA on the simulated positive and null data, respectively, for different sample sizes are shown in Figure 2b and 2d, and was calculated as the fraction of searches returning metafeatures with permutation p-value significant at α=0.05. The TPR was estimated for different number of recovered true positive features (in the true positive datasets) and is summarized in Table 1. The FPR was estimated for different number of returned features (by definition, false positives) in the null datasets, and is summarized in Table 2. CaDrA returned all of the simulated true positive features with 100% TPR for sample sizes larger than *N*=100. CaDrA also yielded a very high mean TPR of > 95% at *N*=100, with the sensitivity dropping to 7.7% only at the smallest sample size of *N*=50. Further, when applied to the null datasets (Fig. 2c), the majority of meta-features returned by CaDrA were correctly deemed as non-significant at α=0.05, with a maximum mean FPR of 7.2% for the lowest sample size analyzed (Fig. 2d). These results suggest that CaDrA requires mid- to large-sized datasets for sufficient sensitivity, while maintaining high specificity at all sample sizes assessed.

**Figure 2.**
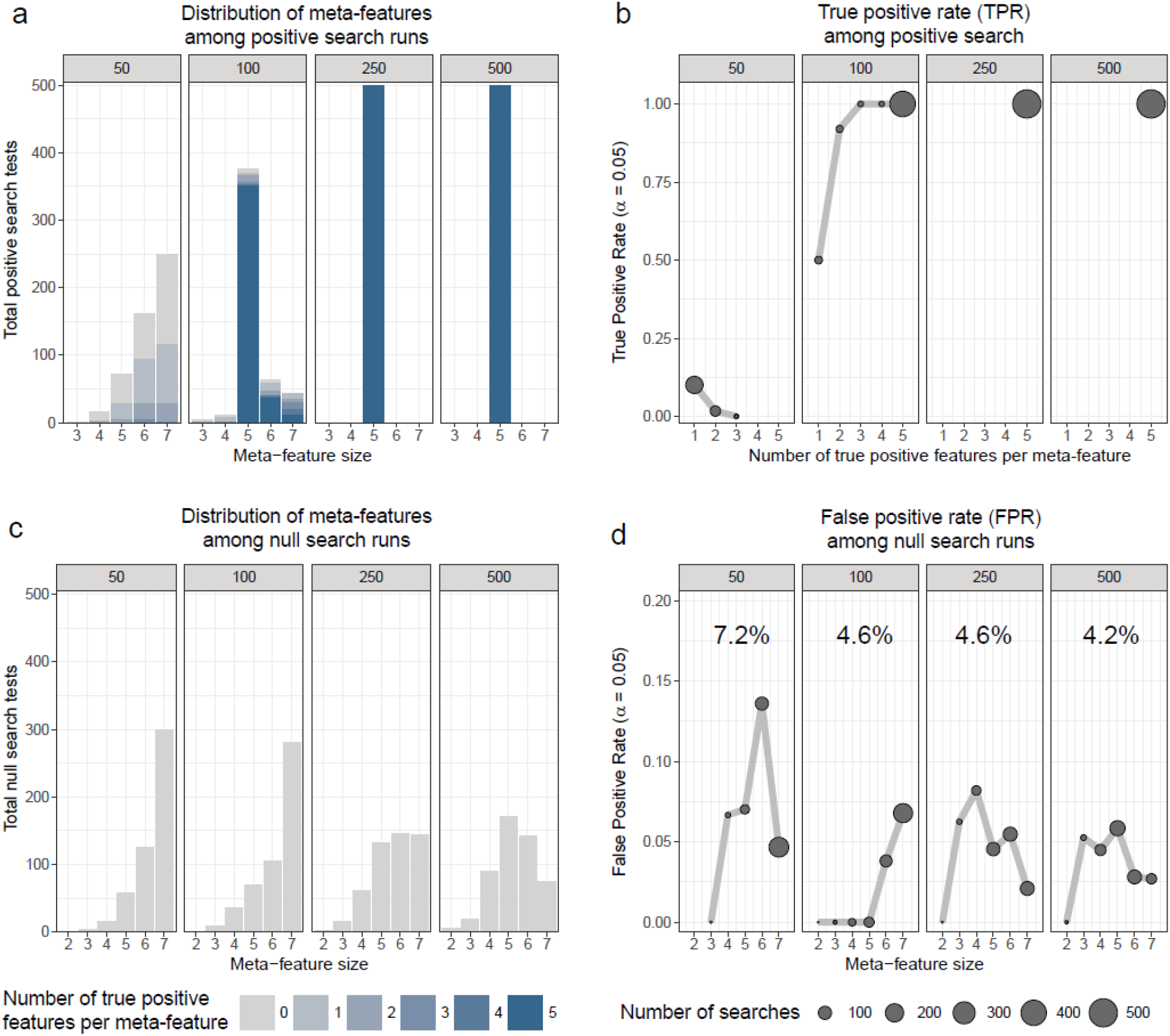
CaDrA performance on simulated data. CaDrA was run on 500 independent simulated datasets containing both positive and null **(a, b)** and only null **(c, d)** features with different sample sizes (gray box above each sub-panel). In each case, the distribution of the number of features per meta-feature (i.e. the meta-feature size) returned by CaDrA is shown **(a, c)** as well as the corresponding number and fraction of searches that yielded significance for α=0.05 **(b, d)**. For example, panel **a** shows that for datasets of size 100, about 375 of the 500 runs returned meta-features of size 5 (i.e., containing 5 features), and in 350 of those runs all five features were true positive (dark blue portion of the bar plot). At sample sizes of 250 and 500, all returned meta-features were of size 5 and all true positives, with statistically significant permutation p-values yielding 100% sensitivity, as indicated in panel b. Panel c shows CaDrA returned metafeatures of different sizes, with majority of these not statistically significant for α=0.05 for all sample sizes considered, as indicated by the weighted average FPRs shown in panel d.

**Table 1.**
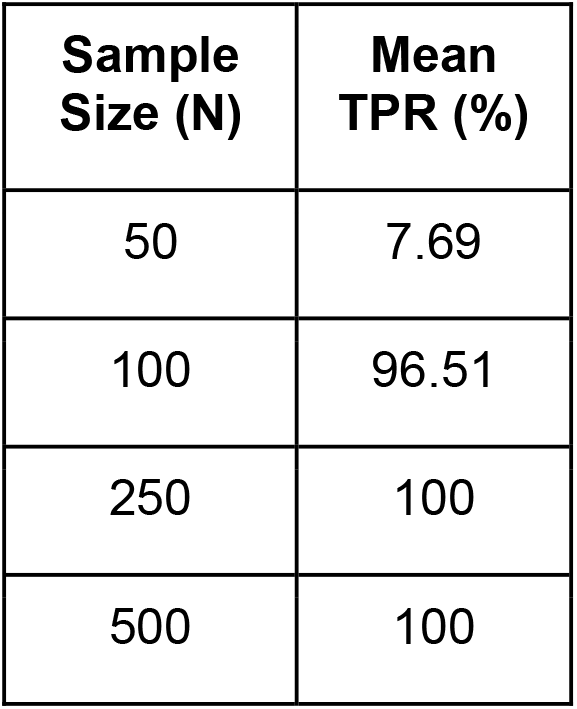
Overall true positive rate (TPR) of CaDrA on true positive simulated data for different sample sizes. Mean percentages shown are a result of weight-averaging TPRs corresponding to different number of true positive features per meta-feature, weighted by the total searches returning such meta-features (see Figure 2b).

| Sample Size (N) | Mean TPR (%) |
| --- | --- |
| 50 | 7.69 |
| 100 | 96.51 |
| 250 | 100 |
| 500 | 100 |

**Table 2.**
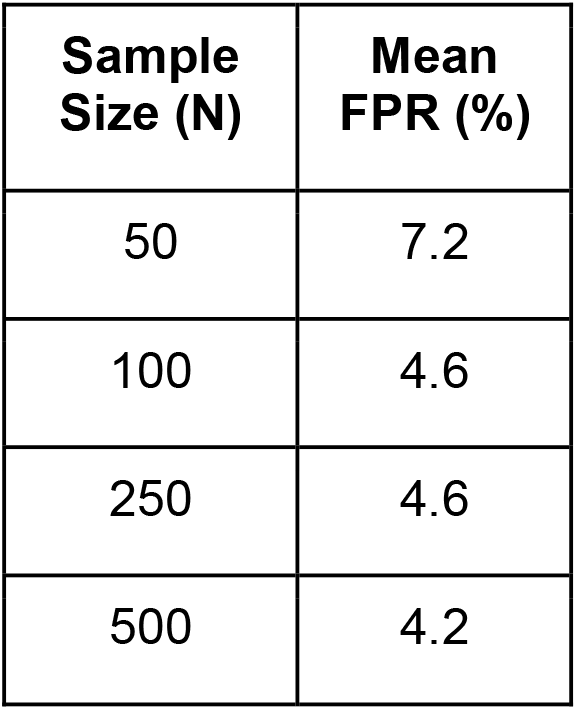
Overall false positive rate (FPR) of CaDrA on null simulated data for different sample sizes. Mean percentages shown are a result of weight-averaging FPRs corresponding to different meta-feature sizes, weighted by the total searches returning such meta-features (see Figure 2d).

### CaDrA identifies known regulators of Ras/Raf/Mek/ERK signaling sensitivity

The mitogen-activated protein kinase (MAPK) kinase (MEKK) / extra-cellular signal-regulated kinase (ERK) pathway is a well-conserved kinase cascade known to play a regulatory role in cell proliferation, differentiation, and survival in response to extracellular signaling [16–18]. Increased MAP/ERK kinase (MEK) activity is a feature of many cancers, and is often triggered by missense mutations in *BRAF* and *NRAS*, two upstream oncogenes and potent regulators of Ras/Raf/Mek/ERK signaling [17,19]. Specifically, the BRAF^*V600E*^ mutation accounts for >90% of BRAF mutations and has been identified as a driver of certain tumors, including melanoma and colorectal cancers, by activating MEK downstream, and is generally found to be mutually exclusive to *NRAS* mutations [19,20]. Small molecules targeting these mutated proteins have been shown to be effective in treating these cancers via inactivation of Ras/Raf/Mek/ERK signaling [1,21–23]. To highlight CaDrA’s ability to recover independent genomic features that may confer hypersensitivity of cancer cells to targeted small molecule treatment, we utilized drug sensitivity profiles for MEK inhibitor AZD6244 [24], along with matched genomic data from CCLE. Specifically, we used per-sample estimates of ‘ActArea’ or area under the fitted dose response curve, a metric that has been shown to accurately capture drug response behavior [25], to rank cell lines from high to low sensitivity, as well as data comprising somatic mutations and SCNAs as the binary feature matrix (see Methods). CaDrA was then run to look for a subset of features associated with increased sensitivity to treatment with AZD6244 (i.e., increased ActArea scores).

The resulting feature set (i.e. meta-feature) is shown in Figure 3. Remarkably, CaDrA selected the BRAF^*V600E*^ and *NRAS* somatic mutations in the first two iterations, respectively. Subsequent iterations identified mutations in *APAF1, TGFBR2* and *AMHR2*, before terminating the search process (P ≤ 0.001). APAF1 is a pro-apoptotic factor and known regulator of cell survival and tumor development [26], the depleted expression of which has been observed in malignant melanoma cell lines and specimens [27]. TGFBR2 and AMHR2 are both type II receptors functioning as part of the transforming growth factor (TGF)/bone morphogenetic protein (BMP) superfamily, together serving as mediators of cellular differentiation, proliferation and survival, and play important roles in directing epithelial-mesenchymal transition (EMT) [28,29]. Notably, MAPK signaling activity can also be regulated by TGF/BMP stimulation [30–32], suggesting that these mutations are potential independent drivers of increased MEK signaling, and hence, of increased sensitivity to treatment with AZD6244. We next extended our analysis of cancer cell line sensitivity profiles to alternative small molecules targeting MEK (PD-0325901), as well as RAF (PLX4720 and RAF265). The meta-features associated with increased sensitivity to each of the four drug treatments assessed are shown in Figure S3 and summarized in Table 3. Importantly, both BRAF^*V600E*^ and NRAS mutations were identified as candidate drivers of sensitivity to MEK inhibition by AZD6244 and PD-0325901. Furthermore, the BRAF^*V600E*^ mutation was returned by CaDrA for all four independent queries, highlighting its association with increased sensitivity to inhibitors targeting the same protein (BRAF) as well as its downstream effector (MEK). Collectively, these results confirm CaDrA’s capability to accurately identify upstream drivers of cellular response to treatment that are both components of independently-linked pathways, as well as part of the same signaling branch, which in turn suggests their role in driving the disease state of interest.

**Figure 3.**
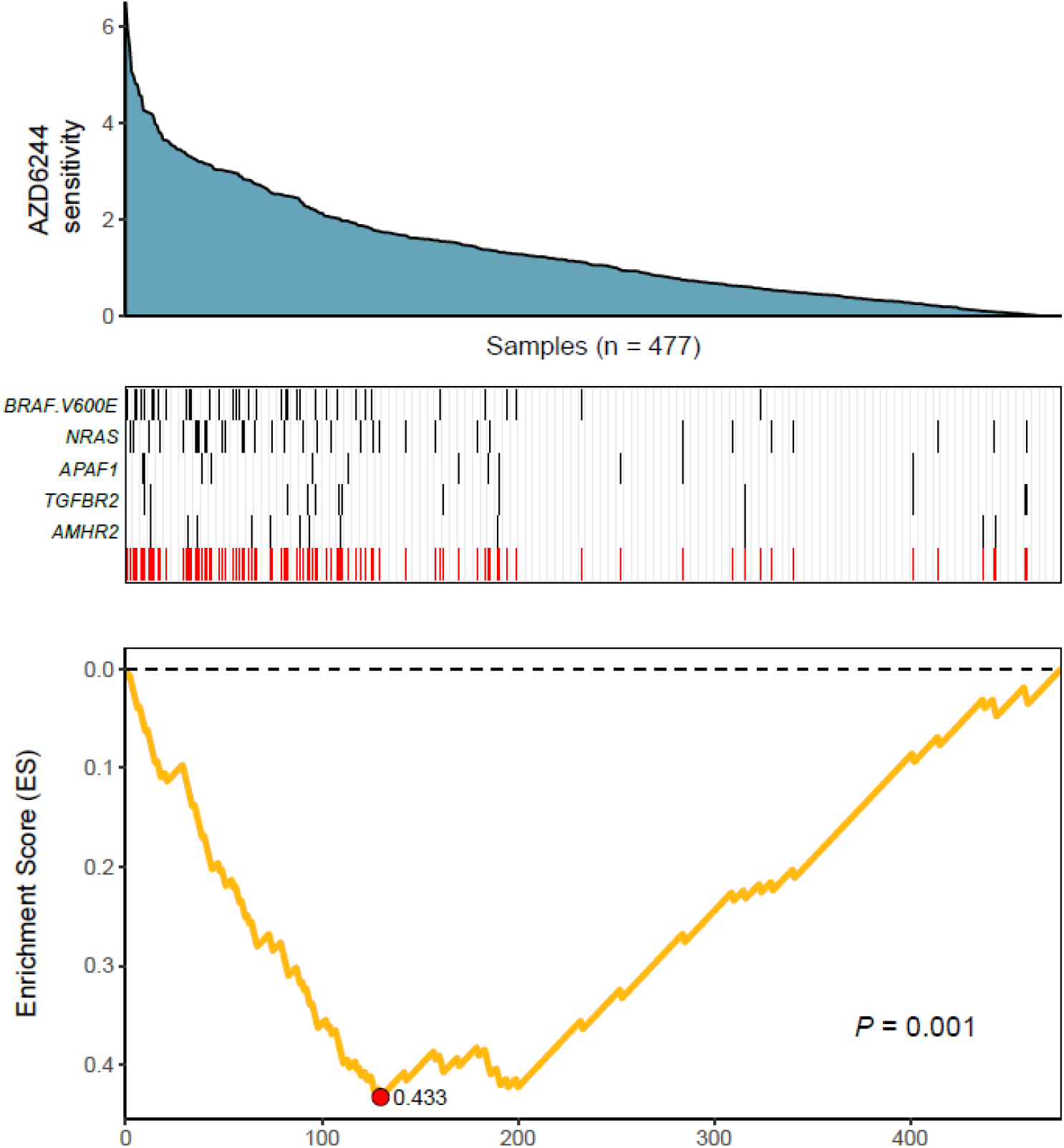
CaDrA identifies mutations in MAPK/ErK signaling genes that contribute to hyper-sensitivity to MEK inhibition *in vitro*. ActArea measurements reflecting sensitivity to MEK inhibitor AZD6244 were used for rank CCLE cell lines (*n*=477). CaDrA was then run to identify sets of genomic features that were most-associated with decreasing ActArea (i.e. increasing sensitivity) scores. Through step-wise search iterations, CaDrA identified somatic mutations in known regulators upstream of MEK, including an activating mutation in *BRAF* (BRAF^*V600E*^) and *NRAS*, as well as those in *APAF1, TGFBR2* and *AMHR2*, before terminating the search process. The resulting metafeature (red track) and its corresponding enrichment score (ES) is shown.

**Table 3.**
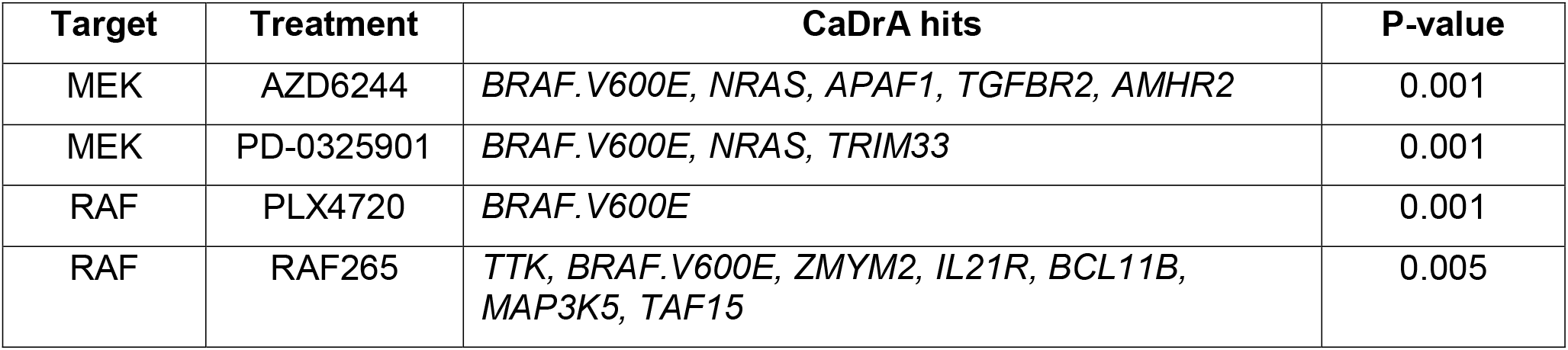
Summary of mutation subsets identified by CaDrA as associated with elevated Mek and Raf inhibition in cancer cell lines. Mutation meta-features identified as associated with increased sensitivity to inhibitors targeting Mek (AZD6244, PD-0325901) and Raf (PLX4720) are shown, along with the corresponding permutation p-value of each search result.

### CaDrA reveals novel drivers of oncogenic YAP/TAZ activity in human breast cancer

The Hippo signaling pathway is a well-characterized highly conserved and tightly-regulated developmental pathway known to play an essential role in cell proliferation and survival [33]. YAP [34] and TAZ [35] serve as central downstream transcriptional effectors of the pathway, with transcriptional activity restricted by Hippo pathway signals that lead to YAP/TAZ nuclear exclusion and degradation. Aberrant nuclear YAP/TAZ localization and activity is associated with a range of cancers, including breast carcinomas (BRCAs) [36–39]. To identify alternative genetic events that can potentially explain the elevated YAP/TAZ activity exhibited in some human breast cancers, we applied CaDrA using genomic data from the TCGA BRCA sample cohort, along with corresponding per-sample estimates of YAP/TAZ activity derived using a gene expression signature of YAP/TAZ knockdown in MDA-MB-231 cells (see Methods). Samples with available RNASeq, somatic mutation and SCNA profiles (*n*=957) were first ranked in decreasing order of their overall YAP/TAZ activity estimates. The ranked binary mutation and SCNA features were then used as input to CaDrA. In the first iteration, CaDrA identified the top scoring genomic feature to be a deletion on chromosomal locus chr5q21.3 (Fig. 4a), harboring tyrosine kinase receptor-encoding gene *EFNA5. EFNA5*, a member of the Eph receptor family, has been hypothesized to function as a tumor suppressor, whose expression has been shown to be reduced in human BRCAs relative to normal epithelial tissue [40]. Advancing to a second iteration, CaDrA then identified an additional deletion of chr20p13 as the next-best feature (Fig. 4a). chr20p13 includes multiple genes (Table S1), including *RBCK1*, whose reduced expression has been shown to be associated with increased tumor cell proliferation and survival, as well as with worse patient survival outcomes in breast cancer [41]. CaDrA then proceeded to identify somatic mutations in the *RELN* gene, before terminating the search process (*P* ≤ 0.001; Fig 4a). Loss of *RELN* expression has indeed been shown to induce cell migration in esophageal carcinoma, and to be associated with poor prognosis in breast cancer [42,43]. To ensure that the derived meta-feature association is not a spurious consequence of correlation with tumor subtype, we tested for the association of YAP/TAZ activity with the meta-feature while controlling for BRCA triple-negative (TN) status. The results confirmed that the positive association between YAP/TAZ activity and the occurrence of these genomic alterations is independent of BRCA patho-histology (Fig. S4; linear regression meta-feature coefficient *P* < 0.0001). Analysis of YAP/TAZ activity based on the same knockdown signature in CCLE BRCA cell lines (*n*=59; Fig. S5a) shows that *RBCK1* and *RELN* display the highest anti-correlation between their gene expression and YAP/TAZ activity (Fig. S5b). In order to assess whether these identified candidates indeed drive the elevated YAP/TAZ activity phenotype, we performed siRNA-mediated knockdown of *RELN* or *RBCK1* in HS578T breast cancer cells, followed by expression quantification of YAP/TAZ canonical targets, which serves as a read-out of nuclear YAP/TAZ activity [44]. HS578T cells which, similar to MDA-MB-231 cells from which the gene signature was derived, are triple-negative BRCA cells but display lower overall YAP/TAZ activity (rank 7/59) compared to the latter (rank 54/59). Importantly, knockdown of either of these candidate drivers in these cells yielded a significant increase in expression levels of YAP/TAZ targets CTGF and CYR61 (FDR < 0.05; two-tailed Student’s t-test), validating the association of their loss of function with increased YAP/TAZ transcriptional activity (Fig. 4b). In summary, unbiased application of CaDrA to the analysis of oncogenic YAP/TAZ activity in primary BRCA samples identified multiple novel candidate drivers of that activity, and our *in vitro* validation confirmed the causal role of the top two candidates, RBCK1 and RELN, in driving that activity, thus providing convincing evidence about the ability of our tool to discover novel drivers.

**Figure 4.**
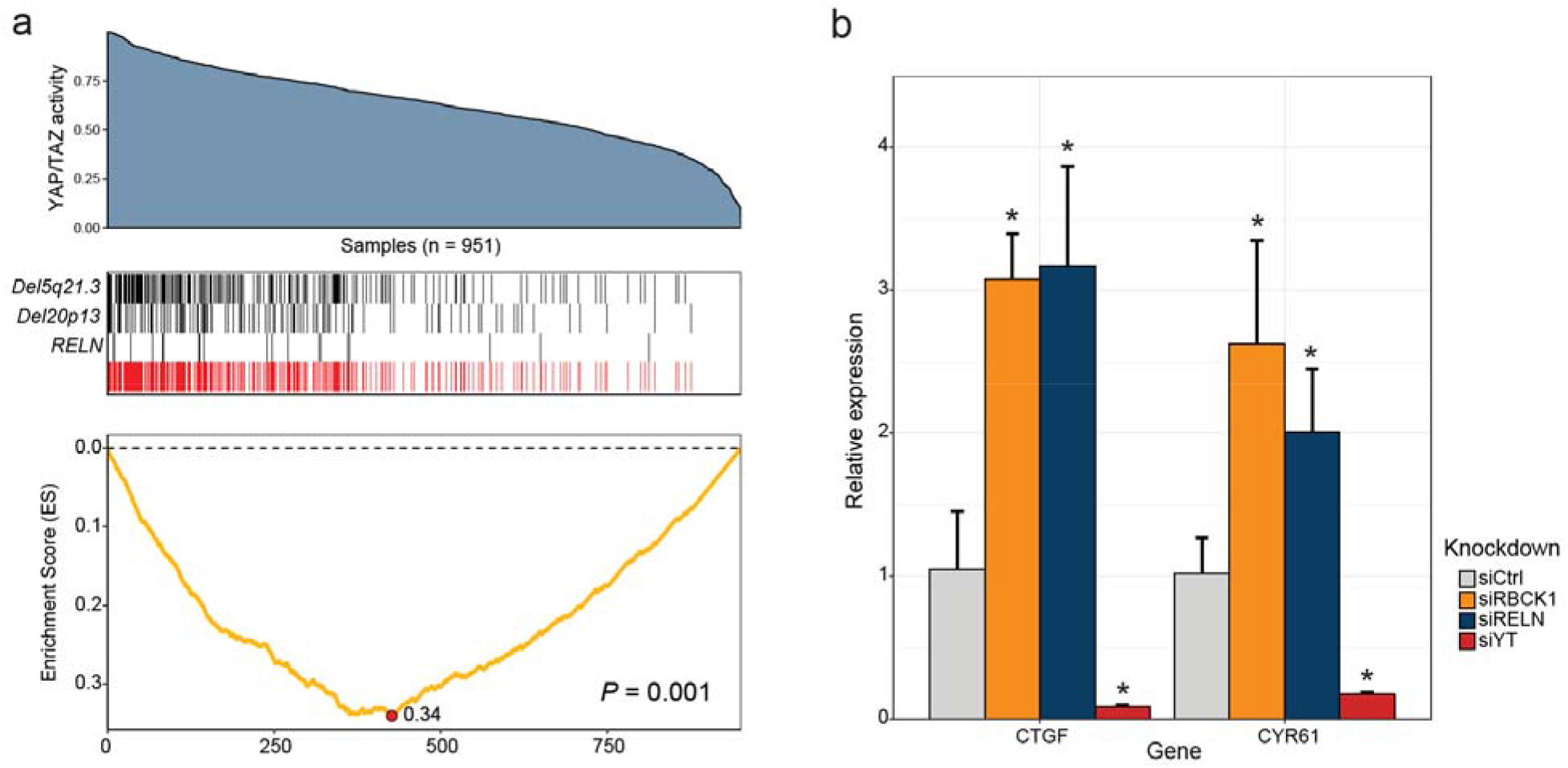
CaDrA identifies novel drivers of oncogenic YAP/TAZ activity in human breast carcinomas. **a**. TCGA BRCA RNASeq data (*n*=951) was projected in the space of a gene signature comprising YAP/TAZ-activating genes to yield a sample stratification based on YAP/TAZ activity estimates (blue area plot; see Methods). CaDrA was then run to look for features associated with elevated YAP/TAZ activity, returning two chromosomal deletions (*Del5q21.3, Del20p13*), and a somatic mutation in *RELN* (black tracks). The union of the three features (red track) and the corresponding running enrichment score (ES) is also shown. b. siRNA-mediated knockdown of 20p13-harboring gene *RBCK1*, and *RELN* in HS578T cells resulted in significant increase in the expression levels of canonical YAP/TAZ targets CTGF and CYR61, as indicated by their relative qRT-PCR expression, confirming the identified CaDrA hits as potential regulators of BRCA-associated YAP/TAZ activity. Ctrl: Scrambled control; YT: YAP/TAZ; * FDR < 0.05; Student’s *t*-test.

## Discussion

Identifying (epi)genetic drivers of molecular readouts is of fundamental importance to determining alternative mechanisms influencing the phenotype in question. Existing methods attempting to extract functionally-relevant sets of genomic alterations associated with a given context either do not support the analysis of data beyond somatic mutations, do not incorporate multiple feature scoring functions and search modes, or do not implement rigorous statistical significance testing of the obtained results. Importantly, a computational framework package bundling all of these features does not exist, and can significantly help identify novel drivers of signature activity.

Here, we presented CaDrA as a tool that determines a subset of queried binary features to be most-associated with a phenotypic signature of interest by specifically exploiting a stepwise heuristic search method based on their union. CaDrA was applied to identify both known and novel genomic drivers of sample signature activity, comprising drug sensitivity and gene set activity estimates, using publically available datasets pertaining to cancer cell lines and primary tumors. Querying CCLE data for features associated with increased sensitivity to Mek/Raf inhibitors, CaDra recovered known driver mutations in oncogenes known to be gate-keepers of MEK pathway activity, including *NRAS* and *BRAF*.

Through our extensive evaluation on simulated data, we were able to highlight CaDrA’s high sensitivity and specificity for mid-to-large sized datasets (*N*>100). Importantly, multi-omic datasets produced by networks such as CCLE and TCGA, also presented in this study, are well above this sample size limit. CaDrA’s specificity was further evident when querying genetic drivers of increased sensitivity to treatment with PLX4720, a potent and selective inhibitor designed to preferentially inhibit active B-Raf protein bearing the V600E allele [45]. In this scenario, the search process correctly identified the BRAF^*V600E*^ mutation as the sole feature associated with elevated sensitivity to treatment, in agreement with the known specificity of the small molecule inhibitor, with the feature association being highly statistically significant. Importantly, we were able to demonstrate the utility of this new framework in the discovery of novel drivers in human breast cancers. Specifically, we asked whether there were genomic alterations associated with elevated activity of Hippo pathway co-activators YAP/TAZ, known to control pro-tumorigenic signals in multiple cancer types [36–38]. The mechanisms contributing to dysregulated YAP/TAZ activity in cancer remain poorly understood. To date, very few genomic alterations have been associated with driving tumorigenic YAP/TAZ activity [46]. Projecting TCGA human BRCA RNASeq data onto a YAP/TAZ-regulated gene expression signature derived from MDA-MB-231 cells, we derived per-sample estimates of YAP/TAZ activity, which was then used as the input ranking variable to CaDrA. CaDrA identified chromosomal deletions of 5q21.3 and 20p13, and mutations in the *RELN* gene as maximally-associated with elevated YAP/TAZ activity based on the union of their occurrences in TCGA samples. Assessment of this identified meta-feature with respect to BRCA TN status confirmed that these genomic alterations are significantly associated with YAP/TAZ activity independent of sample clinical subtype. We wished to validate whether introduction of these identified perturbations would drive a cell towards higher YAP/TAZ activity, in turn, pheno-copying the observed trend in primary BRCAs. Knockdown of select targets, namely RELN and RBCK1, in HS578T BRCA cells exhibiting low YAP/TAZ-activity resulted in a significant increase in the expression of canonical YAP/TAZ targets CTGF and CYR61, confirming their involvement in the regulation of YAP/TAZ-mediated activity. Since CaDrA was designed to identify genomic alterations impacting similar events, we believe these results further emphasize its utility as a tool in identifying and linking novel signaling effectors with a target outcome of interest.

Previously developed methods have indeed been shown to aid in the selection of functionally-relevant genomic features. In particular, REVEALER [13] is an iterative search algorithm that functions in a similar fashion to CaDrA, while specifically seeking only those features that are mutually exclusive given the sample context. We note that a direct and rigorous comparison between our method and REVEALER was not possible given the lack of a formal procedure to estimate statistical significance of results in the latter. We further emphasize that our tool provides a flexible framework capable of incorporating additional feature scoring functions, including the mutual information criterion implemented in REVEALER.

While our evaluations focused on somatic mutations and SCNAs, we note that CaDrA’s search functionality can be applied to additional sequencing readouts, including and not limited to, DNA methylation and microRNA expression, albeit with proper discretization of these continuous features. A joint analysis of these additional data types might provide insight into epigenetic mechanisms that complement the assessed genetic features in driving phenotypic variation. Furthermore, we envision the adoption of CaDrA for the study of germ-line variation as well, thus contributing to move beyond the “one feature at a time” paradigm typical of GWAS studies, although issues of computational efficiency in that problem space will likely become more challenging.

## Conclusions

CaDrA enables one to efficiently identify subsets of genomic features, including somatic mutations and SCNAs, as candidate drivers of a pre-defined phenotypic variable. Given the rapid rise in the availability of multi-omics datasets, as well as an increased need to interrogate targeted molecular readouts within these contexts, we believe that our methodology will accelerate feature prioritization for further follow-up and consideration, in turn aiding in the discovery of potential drivers of the phenotype of interest. Thus, we propose CaDrA as a tool for both targeted hypotheses testing, and novel hypothesis generation.

## Methods

### The CaDrA algorithm

An overview of CaDrA’s workflow is summarized in Figure 1. CaDrA takes as input the sample ranking induced by a sample-specific measurement, a matrix of binary features (1/0 indicating the presence/absence of a given feature in a sample), and a scoring method specification to measure the significance of the concordance between the occurrence of alteration events and the defined sample ranking. The pre-defined sample ranking can be based on quantitative estimates of a gene expression, a signature or pathway activity, or other experimentally-derived measurements. Each row in the matrix of binary features denotes the presence or absence of a somatic alteration (mutation, CNA, or other) in each of the samples in the ranked cohort. The score function is a measure of the *left-skewness* of a binary vector with respect to the sample ranking. The more the occurrences of an alteration are skewed towards higher rankings (i.e., the more the 1’s in the feature vector are skewed towards the left), the higher the score. The scores currently implemented are the Kolmogorov-Smirnov (KS) test, and the Wilcoxon rank-sum test, but additional scoring functions can easily be added.

Given the sample ranking, the matrix of binary features, and the score of choice (KS or Wilcoxon), CaDrA implements a step-wise greedy search: it begins by first selecting the single feature that maximizes the score (Step 1; Fig. 1). It then generates the union (logical OR) of this starting feature with every other remaining feature in the dataset and computes scores for the obtained ‘meta-features’ (Step 2; Fig. 1); it selects a 2^nd^ feature that, added to the first (as a union), maximally increases the score – which will then serve as the new top reference hit (Step 3; Fig. 1). Repeating this process until no further improvement to the cumulative score can be attained, the search output is a set of features (i.e. a meta-feature) whose union has the (local) maximum skewness score with respect to the input sample ranking. The significance of a CaDrA search and its cumulative score are determined by generating an empirical null distribution of scores based on the exact same data and search parameters, but with randomly-permuted sample rankings, providing a permutation p-value per search result.

### CaDrA features

#### Search modes

CaDrA supports multiple search modalities: it allows for the selection of a user-specified feature from which to start the search (rather than selecting the feature with highest score as depicted in Step 1 of Figure 1); alternatively, since the greedy search is not guaranteed to find the global maximum, it also allows for a “top-N” search modality, whereby the search is started from each of the first N features (as measured by their individual skewness scores), and the result of the best search can be determined by selecting the set of features with the best cumulative score over the top-N runs.

#### Visualization of search results

For a given search, CaDrA outputs a set of features (meta-feature), which can be visualized as a ‘meta-plot’. This includes (panels from top to bottom): an area plot of the sample-specific measurements used to obtain the sample ranks; a color-coded matrix of all features in the meta-feature (in the step-wise order that they were added), one feature per row, with the corresponding union of the metafeature (red) last; and a corresponding enrichment score (ES) plot below. Additionally, top-N search results can be visualized for overlapping features to evaluate robustness across different search starting points.

#### Parallelization support

The generation of the empirical null distribution for significance testing is typically done for ≥ 500 iterations (i.e. permuted sample ranks). In order to speed up this potentially time-consuming task, CaDrA supports exploiting parallel computing with the help of the parallel R package functionality, should multiple compute cores be available to users.

#### Permutation caching

Since the generation of the null distribution used for significance testing is a time-consuming step, and since the null distribution of scores depends solely on the feature dataset and the search parameters specified (scoring method, starting feature versus top-N search mode etc.), and not on the input sample ranking, we can implement cacheing of the null distribution corresponding to each dataset and search parameters. When submitting multiple subsequent queries (each with its own sample ranking) that utilize the same dataset and search criteria, CaDrA can then fetch the corresponding cached null distribution to generate permutation p-values almost instantaneously, avoiding the need for repetitive computation, thus significantly reducing overall query run time.

### Data availability and processing

CaDrA is freely available for download and use as a documented R package under the git repository https://github.com/montilab/CaDrA, and will further be deposited and maintained for future use under Bioconductor, including complete code, example use-cases and results pertaining to this manuscript.

RNASeq, DNA copy number (GISTIC2) and mutation data for the TCGA BRCA cohort was downloaded using Firehose v0.4.3, corresponding to the Feb 4^th^ 2015 (RNASeq) and the Jan 28^th^ 2016 Firehose release (SCNA and somatic mutations). Somatic mutation data was processed at the gene level by assigning either 1 or 0 based on the presence or absence of any given mutation in that gene, respectively (excluding synonymous mutations). RNASeq version 2 data corresponding to Level 3 RSEM-normalized gene expression values were used. CCLE genomic data were downloaded from https://portals.broadinstitute.org/ccle and processed as previously described [13]. Somatic mutation binary calls per gene were used as is, and SCNA data was processed using GISTIC2 [47] with all default parameters barring the confidence level, which was set to 99%. ActArea estimates pertaining to drug treatment sensitivity across CCLE samples was used as previously described [1].

In all cases presented, SCNA and somatic mutation data were jointly analyzed as a single input dataset to CaDrA, thereby including samples for which both data were available. All input data to CaDrA were further pre-filtered so as to exclude alteration frequencies below 3% and above 60% to reduce feature sparsity and redundancy, respectively, across samples (CaDrA’s default feature pre-filtering settings).

### Simulated data generation

To evaluate both the sensitivity and specificity of CaDrA, we generated simulated data to represent cases where there was a mix of left-skewed (“true positive”) and randomly distributed (“null”) features, as well as cases where there were only null features. The left-skewness of a feature is a measure of its association with the sample ranking, since samples are sorted from left (high rank) to right (low rank). The design and parameter specification of the simulated data matrix is shown in Figure S1. Each feature/row is a binary (0/1) vector, with 1 (0) in the *i*^th^ position denoting the occurrence (nonoccurrence) of the genetic event (e.g., SCNA or mutation) in the *i*^th^ sample. This simulation of binary features relies on the following parameters:

*N*: Dataset sample size (number of columns in the matrix)
*n*: Total number of features in the dataset (number of rows in the matrix)
*p*: Number of true positive features generated per dataset (a positive feature is a feature whose distribution of events (i.e. the number of 1’s) is significantly associated with the sample ranking, i.e., left-skewed).
*f*: Left-skew proportion. The proportion of samples that are *cumulatively* left-skewed in the sample ranking.
*λ*: The mean (and variance) of the Poisson distribution from which the number of events in the null features is sampled. This is equal to the number of 1’s per skewed positive feature. A Poisson distribution is used so that we can partially control (through the mean) the number of 1’s in a null feature, which are then uniformly distributed across samples (see description of Null feature generation below).

The resulting simulated binary data matrix will consist of two main types of features: *True Positive (TP) features*: A total of *p* TP features are generated. Events (i.e., 1’s) are assigned to the TP features in a mutually exclusive fashion, with each of these features having (*f×N*) /*p* entries set to 1, with their cumulative OR yielding an N-sized vector with the left-most *f*×*N* entries set to 1’s. For example, if we generate data for 100 samples and 5 positive features, with the left-skew proportion set to 0.5, each non-overlapping feature will have 10 among the 50 left-most entries (columns) set to 1, such that the union (logical OR) of the 5 features will have 1’s in the first 50 entries.

#### Null features

Null features are generated for a total of (*n* - *p*) features. To generate these features, we sample the number of 1’s per null feature based on a Poisson distribution with mean parameter λ = (*f*×*N*) / *p*. In this fashion, the number of 1’s in the null features will have a distribution centered on the corresponding number for the TP features. For instance, if we generate data for 100 samples and 5 TP features with left-skew proportion *f*=0.5, then each of the TP features will have ten 1’s, and each of the remaining 995 null features will have a number of 1’s sampled from Poisson (λ=10), uniformly distributed over the *N* samples.

A schematic representation of this data, along with the parameters that define its composition is shown in Figure S1.

### Evaluation of CaDrA performance on simulated data

Evaluation of CaDrA performance was performed considering two main scenarios: a) True positive datasets: Data containing both true positive and null features (where the sensitivity of CaDrA is tested); and b) Null datasets: Data containing only null features (where the specificity of CaDrA is tested), with the following parameter specifications for data generation:

*N* = {50, 100, 250 and 500}
*n* = 1000
*p* = 5
*f* = 0.5

CaDrA was run using default input parameters, returning a meta-feature which had the best score, along with a permutation p-value based on the empirical null search distribution (Fig. S2). These results were then used to determine performance estimates for different sample sizes, composition (i.e. distribution of TP versus null features per returned meta-feature), size (i.e. the number of features within the returned meta-feature) and statistical significance of the returned meta-features.

### YAP/TAZ signature projection and assessment in TCGA BRCAs

A signature comprising YAP/TAZ-activating genes (*n*=717) in MDA-MB-231 cells was obtained based on a previous study [48]. The TCGA BRCA RNASeq data (*n*=1,186 samples) was projected onto the signature genes and per-sample estimates of YAP/TAZ activity were derived using ASSIGN [49], which was then used as a continuous ranking variable with CaDrA. The association of YAP/TAZ activity with the CaDrA-derived meta-feature, and with BRCA subtype (i.e. triple-negative status) was determined using a linear regression model.

### Cell culture, siRNA knockdown and qRT-PCR

HS578T BRCA cells were purchased from ATCC and cultured using media and conditions suggested by ATCC. For RNA interference, cells were transfected using RNAiMAX (thermofisher) with control siRNA (Qiagen, 1027310) or an equal molar mixture of siRNA targeting RELN (Sigma), RBCK1 (Sigma), or TAZ and YAP [50]. 48 hours post transfection, RNA was extracted from cells using RNeasy kit (Qiagen) and the synthesis of cDNA was performed as previously described [50]. Quantitative real-time PCR (qRT-PCR) was performed using Taqman Universal master mix II (thermofisher) and measured on ViiA 7 real-time PCR system. Taqman probes used included those recognizing CTGF (thermofisher Hs00170014_m1), CYR61 (thermofisher Hs00155479_m1), RELN (thermofisher Hs01022646_m1), RBCK1 (thermofisher Hs00934608_m1), WWTR1 (thermofisher Hs01086149_m1), and YAP (thermofisher Hs00902712_g1) and GAPDH (thermofisher 4326317E). Expression levels of each gene were calculated using the ΔΔCt method and normalized to GAPDH. Knockdown efficiency of YAP, TAZ, RELN and RBCK1 was verified for each experiment. Mean transcriptional knockdown of YAP, TAZ and RBCK in HS578T cells was > 80%. Basal RELN levels in HS578T cells were low, and relative knockdown in these cells was 28.3% (±14.1). Data from qRT-PCR experiments are shown as mean ± S.D., with each knockdown compared with respect to the scrambled siRNA control (siCtl) using an unpaired, two-tailed Student's *t*-test.

### CaDrA search parameters

For evaluation using genomic data, CaDrA was run in the top-N mode using the default of N=7, choosing the best resulting meta-feature. (see Methods; CaDrA features: Search modes). For evaluation of simulated data, only the top-scoring feature was considered as a starting feature per search run (i.e. N=1). All other default input search parameters were used for all cases presented.

## List of abbreviations

CaDrA: : Candidate Driver Analysis
TCGA: : The Cancer Genome Atlas
CCLE: : Cancer Cell Line Encyclopedia
BRCA: : Breast Carcinomas
TN: : Triple-negative
TPR: : True Positive Rate
FPR: : False Positive Rate
FDR: : False Discovery Rate
qRT-PCR: : Quantitative Real-Time Polymerase Chain Reaction

## Declarations

### Ethics approval and consent to participate

Not applicable.

### Consent for publication

Not applicable.

### Availability of data and material

The datasets generated and/or analyzed during the current study are available in the TCGA repository (https://tcga-data.nci.nih.gov/docs/publications/tcga), and CCLE repository (https://portals.broadinstitute.org/ccle), and are available from the corresponding author on reasonable request.

### Competing interests

The authors declare that they have no competing interests.

### Funding

This work was supported by National Institutes of Health NIDCR fellowship F31 DE025536 (VKK), CDMRP W81XWH-14-1-0336 (XV) and the Dahod breast cancer research program at Boston University School of Medicine (XV and SM). The funding sources played no role in the design of the study and collection, analysis, and interpretation of data and in the writing of this manuscript.

### Author’s contributions

VKK developed the R package. VKK, SM, PS and XV wrote the manuscript. JGK performed the siRNA and qRT-PCR experiments. LZ assisted in obtaining the gene expression signature for TCGA data projection. PS helped in the evaluation of CaDrA on simulated data. SM and VKK designed the CaDrA framework and features, and interpreted the results. XV designed the experimental validation of novel candidate drivers, and interpreted the results thereof. All authors read and approved the final manuscript.

## Acknowledgements

We would like to thank Joshua Klein for making suggestions towards the implementation of specific package features. We further acknowledge dbGap for granting access to the TCGA data (phs000178.v9.p8).

